# Sequential *in vivo* labeling of insulin secretory granule pools in INS-SNAP transgenic pigs

**DOI:** 10.1101/2021.04.23.441054

**Authors:** Elisabeth Kemter, Andreas Müller, Martin Neukam, Anna Ivanova, Nikolai Klymiuk, Simone Renner, Kaiyuan Yang, Johannes Broichhagen, Mayuko Kurome, Valerie Zakhartchenko, Barbara Kessler, Klaus-Peter Knoch, Marc Bickle, Barbara Ludwig, Kai Johnsson, Heiko Lickert, Thomas Kurth, Eckhard Wolf, Michele Solimena

## Abstract

β-cells produce, store and secrete insulin upon elevated blood glucose levels. Insulin secretion is a highly regulated process. The probability for insulin secretory granules to undergo fusion with the plasma membrane or being degraded is correlated with their age. However, the molecular features and stimuli connected to this behavior have not yet been fully understood. Furthermore, our understanding of β-cell function is mostly derived from studies of *ex vivo* isolated islets and/or rodent models. To overcome this translational gap and study insulin secretory granule turnover *in vivo*, we have generated a transgenic pig model with the SNAP-tag fused to insulin. We demonstrate the correct targeting and processing of the tagged insulin and normal glycemic control of the pig model. Furthermore, we show specific single- and dual-color granular labeling of *in vivo* labeled pig pancreas. This model may provide unprecedented insights into the *in vivo* insulin secretory granule behavior in an animal close to humans.

## Introduction

*β*-cell dysfunction is a key contributor to type 2 diabetes mellitus (T2DM) (Solimena et al., 2018; Saeedi et al., 2019) starting in the early onset of the disease (Cohrs et al., 2020). Each *β*-cell contains several thousand insulin secretory granules (SGs) (Fava et al., 2012; Müller et al., 2020). However, only a small percentage of insulin SGs undergoes exocytosis upon glucose stimulation (Rorsman and Renström, 2003). Insulin is secreted in two phases: a rapid first and a sustained second phase (Cerasi and Luft, 1967; Curry et al., 1968; Jaffredo et al., 2021). On the level of insulin SGs our understanding of insulin secretion has been shaped by two basic concepts: The recruitment of SG pools defined by their spatial confinement in the cell and the higher probability of young SGs for exocytosis. In the first model the so-called readily-releasable pool consists of SGs that are already docked with the plasma membrane and are released immediately upon glucose stimulation, thereby creating the first rapid phase of insulin secretion (Barg et al., 2002). The second prolonged phase is then caused by the recruitment of SGs from the reserve pool which is located deeper inside the *β*-cell (Rorsman and Renström, 2003). The detailed properties of SGs of the different pools have been debated and refined recently (Ohara-Imaizumi et al., 2007). Additionally, data obtained by radio-labelling experiments suggest that young insulin SGs are preferentially secreted (Schatz et al., 1975; Halban, 1982). A method that allows for the visualization of age-defined pools of the desired protein is to fuse it with the SNAP-tag, a 20 kDa protein tag that reacts covalently in a bioorthogonal manner with fluorescent benzylguanine-fused substrates in living cells and organisms (Keppler et al., 2003). By using a pulse-chase labelling approach to track SGs containing an insulin-SNAP chimera, we could confirm the preferential exocytosis of young SGs and could also show the preferential intracellular degradation of old SGs (Ivanova et al., 2013; Müller et al., 2017b,a). Furthermore, we found in insulinoma INS-1 cells that a pool of young SGs travels fast on microtubules, while this property is lost for old SGs (Hoboth et al., 2015). Young insulin SGs additionally have a more acidic lumenal pH compared to old ones (Neukam et al., 2017). Addressing the heterogeneity of insulin SGs and their differential reaction to stimuli and pharmaceutical intervention poses novel possibilities for treatment of T2DM. Genetically modified mouse models have been the method of choice to investigate intracellular signaling as well as metabolism in diabetes research. Recently, transgenic pigs have been made available that allow for conducting β-cell research in a context closer to humans (Wolf et al., 2014). Here, we describe the generation and characterization of a transgenic pig model with the SNAP-tag fused to insulin called SOFIA (Study OF Insulin granule Aging) pig. We demonstrate the correct targeting and processing of insulin-SNAP to insulin SGs. Finally, we show successful *in vivo* labeling with one and two SNAP-substrates staining pancreatic islets and distinct insulin SG pools. In summary, our pig model is a valuable system enabling imaging-based investigation of insulin SG turnover in a large living mammal.

## Results

### Generation of a transgenic pig model (*SOFIA* pig) expressing Insulin-SNAP

To monitor the intracellular trafficking and turnover of the insulin SGs, we designed an expression vector of porcine insulin gene containing the porcine *INS* core promoter and genomic fragments from the porcine INS gene for proper splicing and polyadenylation of the transgene (Fig. 1A). The SNAP-tag was cloned in frame at the 3’-end of the INS coding sequence. After transfection of the vector into porcine kidney cells, their positive selection and depletion of the floxed neoR cassette, we used a mixed population of genetically modified cell clones in somatic cell nuclear transfer (SCNT) experiments and transferred cloned embryos to estrus-synchronized gilts. In total, eleven *INS-SNAP* transgenic (*SOFIA*) founder piglets with ten different integration patterns were obtained (Fig. 1B). The highest transgene expression was detected specifically in β-cells of founder 1817 (Supp. Fig. 1A). This founder pig had two segregating transgene integration sites: one with full Cre-mediated deletion of the floxed neo cassette exhibiting medium transgene expression levels, and another with two *INS-SNAP* copies, one with and the other without neo deletion. The latter integration site resulted in the highest *INS-SNAP* expression (Fig. 1B, Supp. Fig. 1A) and was used to set up a pig line for further experiments. Immunostaining and confocal microscopy for the SNAP-tag and insulin showed colocalization of SNAP with insulin in the islets of Langerhans of transgenic offspring, whereas *wild type* (*WT*) pig pancreas was negative for SNAP (Fig. 1C, Supp. Fig. 2). To determine the transgene integration site(s), we performed targeted locus amplification (TLA) with subsequent next generation sequencing (NGS). The results demonstrate that the transgene is integrated at a single locus on chromosome 11 in a non-coding genomic region with a nearest distance of >0.4 Mb to the next coding gene (Supp. Fig. 1B). In detail, integration of the transgene resulted in the duplication of the genomic locus chr11:56,928,515-56,937,394. These duplicated regions flank the transgene, which is present at either of two possible orientations (Supp. Fig. 1C). As observed by Southern Blot (Supp. Fig. 1A), the sequencing confirmed the presence of one *INS-SNAP* cassette with, and another without, the floxed neo-cassette (Supp. Fig. 1C). Taken together, these data show the successful integration of the *INS-SNAP* transgene in the porcine genome in a single locus on chromosome 11.

**Fig. 1.**
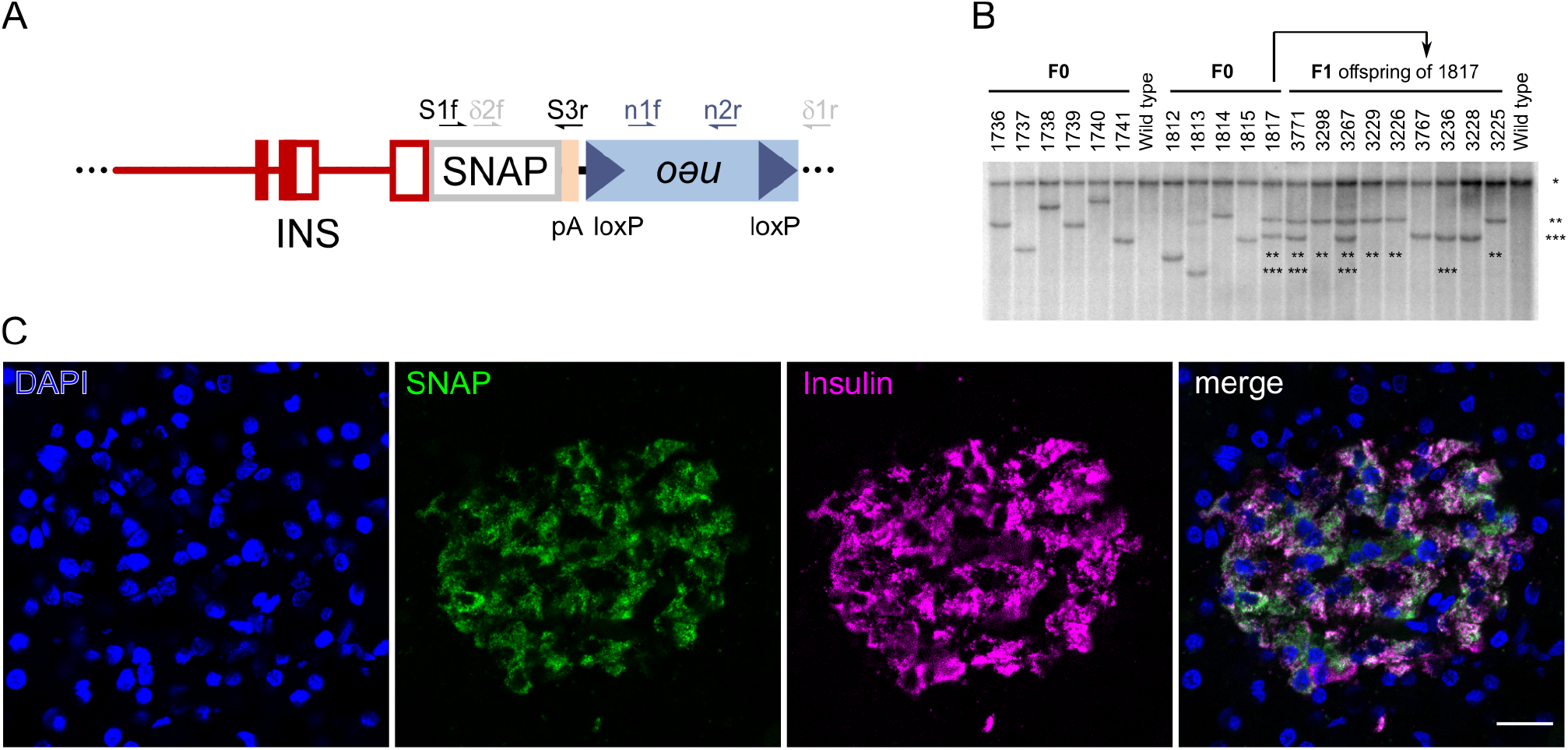
Generation and characterisation of *SOFIA* founder pigs and offspring. (A) *β*-cell-specific *INS-SNAP* expression vector with 1.3 kb upstream regions, exon 1-3 and intron 1 of the porcine INS gene, in frame SNAP-tag sequence and a poly-adenylation (pA) cassette of the bovine growth hormone (*GH*) gene, linked to a floxed neomycin resistance cassette. Site of primers used for genotyping are indicated. (B) Southern blot analysis for evaluation of integration pattern. Integration patterns of eleven F0 founder pigs and nine F1-offspring of founder 1817 are shown. gDNA was digested with the ‘null cutter’ restriction endonuclease enzyme Ndel. *, the band representing the endogenous INS promoter; **, neo/*δ*neo transgene integration site with two integrants where neo cassette was deleted only in one *INS-SNAP* integrant; ***, *δ*neo transgene integration site. (C) Immunofluorescence labeling against SNAP-tag and insulin in transgenic offspring in the *SOFIA* F2 generation. Scale bar: 20 μm.

### Characterization of Insulin-SNAP expression and targeting

Next, we characterized the expression levels and localization of INS-SNAP on the cellular level by real-time quantitative PCR (RT-qPCR), immunoblotting and microscopy. To this aim, we isolated mRNA from total pancreas for RT-qPCR. As expected, the transgene was not present in WT animals (Fig. 2A), but comparable to γ-*tubulin* expression level in *SOFIA* pig pancreata. Endogenous insulin, however, in both *SOFIA*, as well as *WT* pigs, was ~600-fold higher than the transgene or γ-*tubulin*. Further, we crossed *SOFIA* pigs with *INS-eGFP* animals (Kemter et al., 2017) to allow for fluorescence activated cell sorting (FACS) of β-cells from pancreas. The sorted cells were then used for immunoblotting. Both *SOFIA/INS-eGFP* and *INS-eGFP* animals showed comparable levels of 9-12 kDa proinsulin and 6 kDa insulin in FACS sorted β-cells (Fig. 2B). When probing for the SNAP tag, however, only β-cells of the *SOFIA* pig showed a robust signal, shifted about ~20 kDa, resulting from the additional 19.4 kDa SNAP tag. Finally, we applied correlative light and electron microscopy (CLEM) to fixed pancreas tissue obtained from adult *SOFIA* pigs. Immuno-labeling with a primary antibody against SNAP-tag followed by a secondary Alexa 488-coupled antibody and protein-A-gold 10 nm showed granular subcellular staining (Fig. 2C upper panel left and middle). Overlay with the electron microscopy (EM) image allowed for the identification of the fluorescent signal within insulin SGs in islets of Langerhans (Fig. 2C upper panel right). This fluorescent signal further correlated with protein-A-gold labeling. The ultrastructure of SNAP-positive insulin SGs appeared to be normal with insulin SGs containing one or more insulin crystals surrounded by a translucent halo (Fig. 2C lower panel). Overall, the ultrastructure of the β-cells of *SOFIA* pigs was comparable to that of pigs not expressing insulin-SNAP and to published electron microscopy data of *WT* pigs (Vantyghem et al., 1996; Lukinius and Korsgren, 2001) with normal appearance of insulin SGs, mitochondria and endoplasmic reticulum without any signs of stress or structural alterations (Supp. Fig. 3). Furthermore, all insulin SGs in SNAP-positive β-cells contained SNAP-tagged insulin (Fig. 2C lower panel right).

**Fig. 2.**
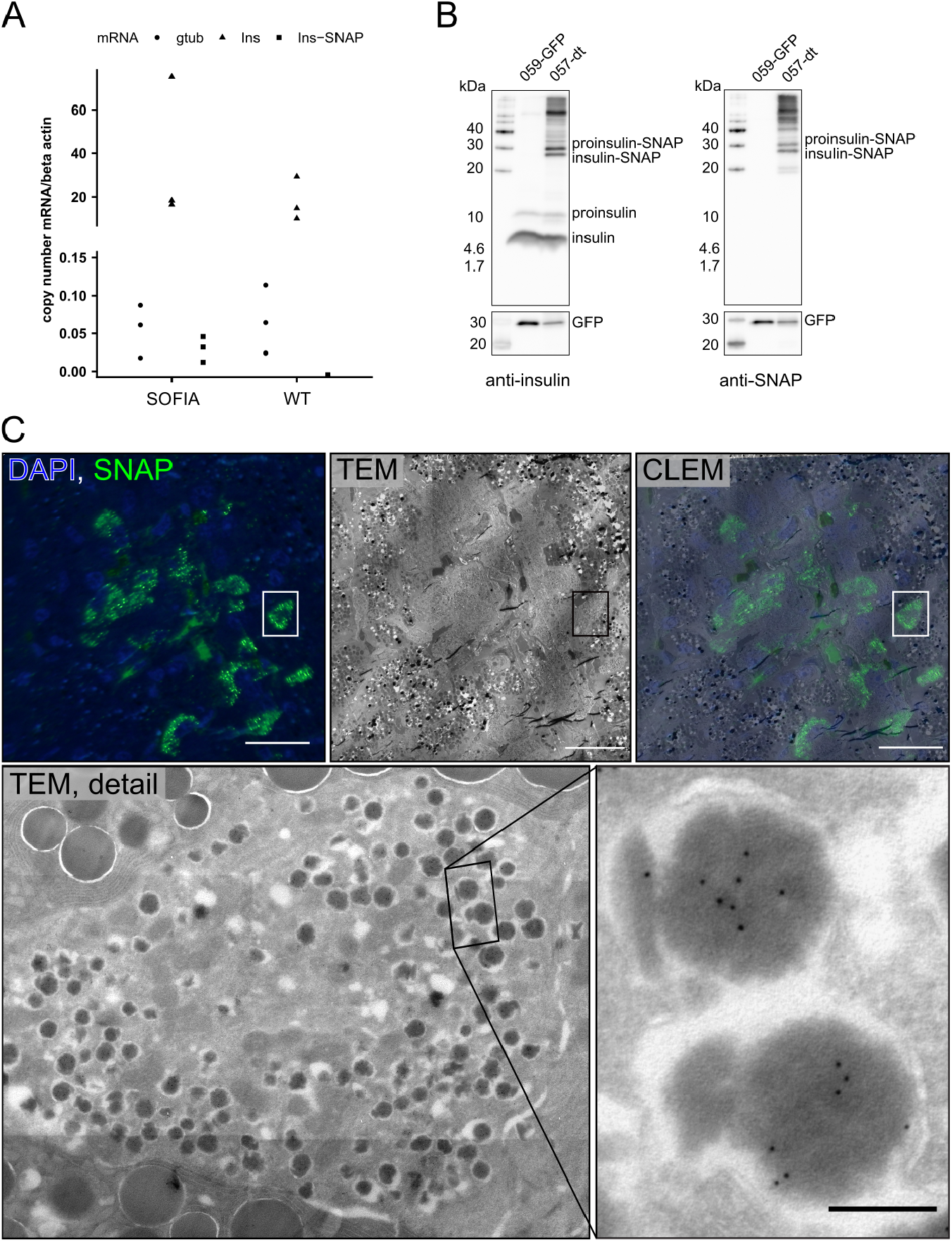
Expression and targeting insulin-SNAP to insulin SGs in *SOFIA* pig islets. (A) qPCR of *SOFIA* pig and WT pig pancreas. (B) Western blots against insulin and SNAP on FACS sorted *GFP-SOFIA* or *GFP-WT β*-cells. 059-GFP: *β*-cells from *INS-eGFP* animal; 057-dt: beta cells from *SOFIA/INS-eGFP* animal. (C) CLEM of *SOFIA* pig pancreas. Fluorescence image shows anti-SNAP labeling (green) with SNAP+ *β*-cells and DAPI (blue). Corresponding TEM image shows the pancreatic islet surrounded by exocrine tissue. CLEM overlay shows SNAP+ signal within *β*-cells (scale bars: 20 μm). TEM detail shows SNAP+ *β*-cell with inset showing immunogold labeling against insulin SNAP (10 nm gold), scale bar:200 nm.

### *SOFIA* pigs have normal glucose tolerance

To assess metabolic control of insulin secretion on the organism level, we performed *in vivo* glucose tolerance tests (IVGTT) in *SOFIA* and *WT* pigs (Fig. 3). After administration, glucose was rapidly cleared from the blood of either pig strains without any obvious difference (Fig. 3A, B). Simultaneously, plasma insulin levels were comparable between the two groups without any significant difference (Fig. 3C, D). Accordingly, further parameters, such as the quotient of insulin and glucose, the insulin sensitivity index and the acute insulin response were not significantly altered in *SOFIA* pigs in comparison to *WT* pigs (Fig. 3E, G). In conclusion, these data indicate that the insertion of the INS-SNAP transgene in this *SOFIA* pig strain does not interfere with glucose homeostasis *in vivo*.

**Fig. 3.**
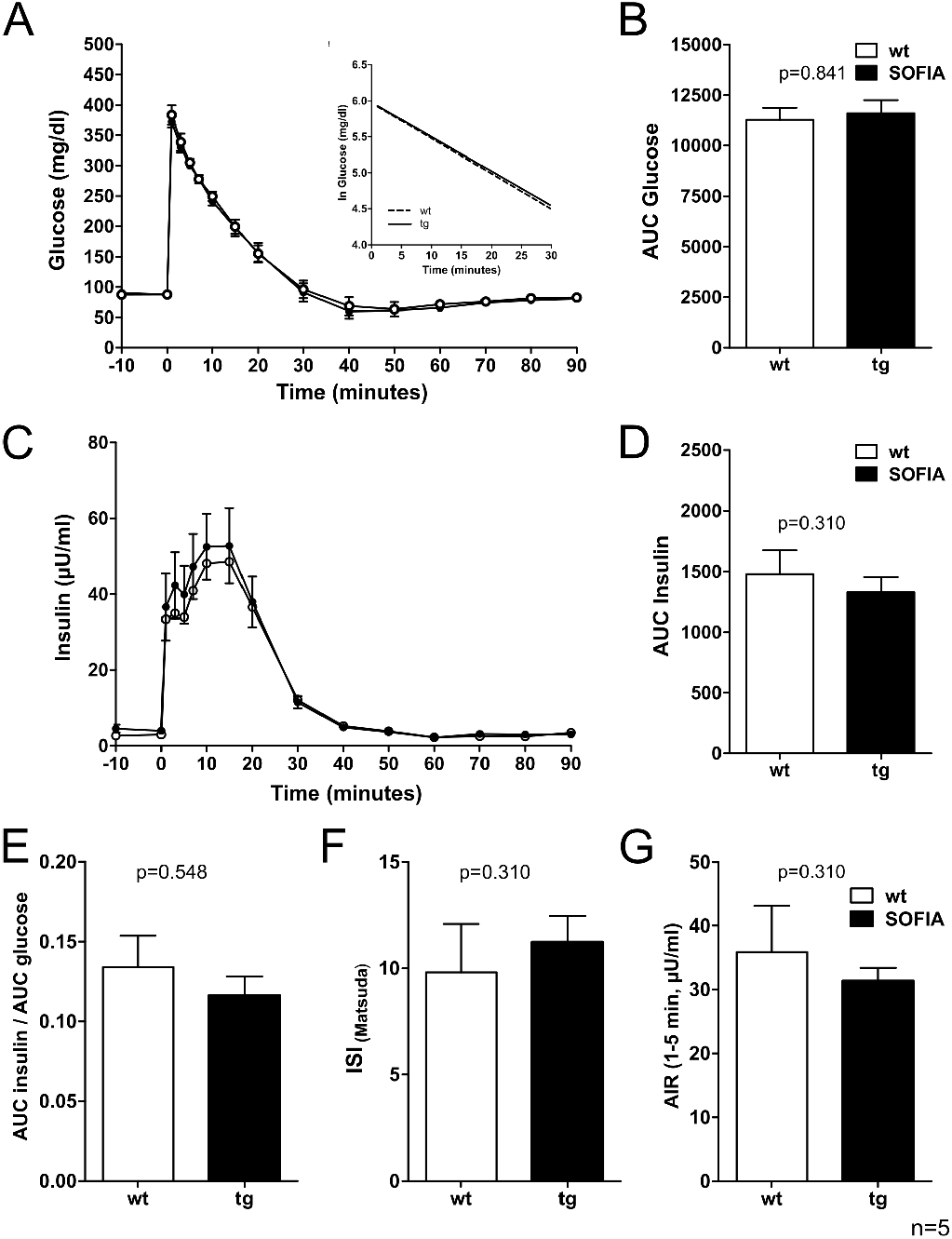
IVGTT and related indices of 15-16-week-old *SOFIA* pigs and *WT* controls. (A) Plasma glucose levels; Insert: glucose elimination rate; (B) Area under the curve for glucose (AUC glucose); (C) Plasma insulin levels and (D) AUC insulin; (E) Quotient of AUC insulin and AUC glucose; (F) Insulin sensitivity index according to Matsuda (ISI Matsuda); (G) Acute insulin response (AIR). Data are represented as means ± SEM.

### SNAP-labeling of insulin SGs *in vivo*

Next, we investigated the functionality and suitability of the SNAP tag for age-dependent labeling of insulin SGs in living pigs. To this aim, we injected 0.6 - 2 μmol/100 kg body weight (BW) of fluorescent benzylguanine (BG) SNAP substrates intravenously in *SOFIA* and *WT* pigs. Injection of BG coupled with tetramethylrhodamine (BG-TMR) followed by sacrifice of the animal and fixation of the pancreas resulted in a clearly detectable TMR signal in cryosections of the pancreas (Fig. 4A). The signal was restricted to the pancreatic islets and was co-localized with the insulin antibody staining (Supp. Fig. 4). Although a fluorescent TMR fluorescent signal could be selectively imaged after *in vivo* application of 0.6 μmol TMR per 100 kg BW, we decided to apply 1.8 - 2 μmol SNAP substrate per 100 kg BW for in vivo SNAP labeling to obtain a more robust imaging signal. For comparison, *SOFIA* mice received 15 nmol SNAP substrate per mouse (15 nmol/25 g) (Ivanova et al., 2013), i.e. 60 μmol/100 kg body weight. Hence, for the *in vivo* labeling of *SOFIA* pigs we applied 30-fold less SNAP substrate per kg of body weight. In order to demonstrate the suitability of the approach for labeling distinct insulin SG pools we performed a dual-color labeling experiment with a first application of BG-TMR followed by BG-silicon rhodamine (SiR) after 15 hours and autopsy and fixation of the pancreas 2 hours later. Again, we could detect the red and far-red fluorescence signals of both SNAP substrates in the islets and detect granular labeling at high magnification (Fig. 4B). Furthermore, there was co-localization of TMR+ and SiR+ insulin SGs as well as the occurrence of SGs labeled with only a single substrate indicating the segregation of SG pools over time. Taken together, these data show that INS-SNAP can be efficiently labeled *in vivo* by injection of fluorescently modified SNAP substrates and further shows suitability for age-dependent labeling of distinct SG pools.

**Fig. 4.**
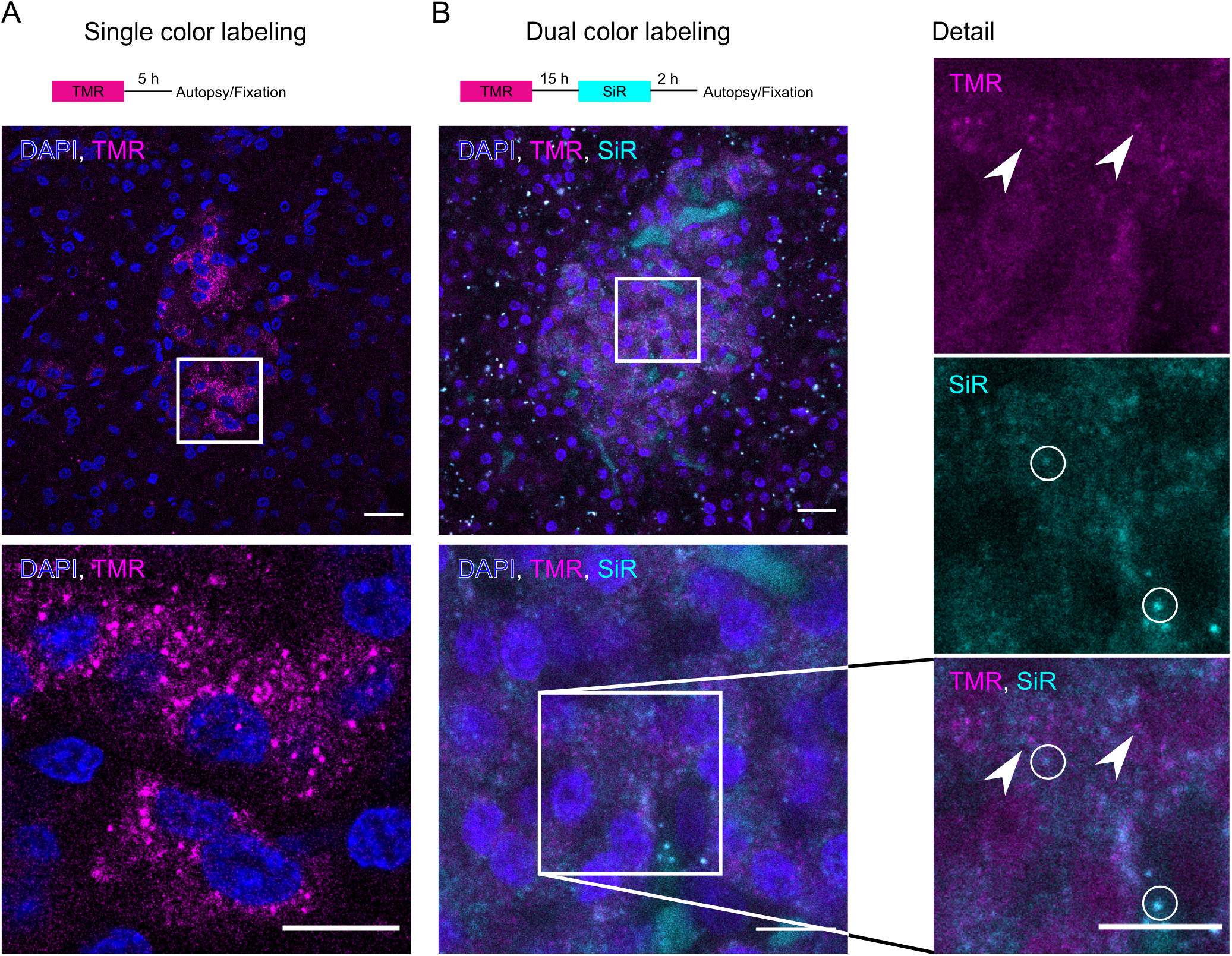
*in vivo* labeling of *SOFIA* pigs. (A) Scheme for single color BG-TMR labeling. Below is a confocal microscopy image of a cryo section showing TMR fluorescence in the islets of Langerhans (scale bar: 20 μm). The magnified view shows granular TMR fluorescence (magenta) and nuclei (blue) (scale bar: 10 μm). (B) Scheme for dual color labeling with BG-TMR and BG-SiR. Below is a confocal image of a cryo section of a *SOFIA* pig showing TMR+ and SiR+ granular staining with nuclei (blue). Detailed views show the magnified boxed area with split TMR and SiR channels. Arrowheads point to exclusively TMR+ SGs and circles show SiR positive SGs. (scale bar: 10 μm).

## Discussion and Outlook

In the present study we have successfully generated and characterized a transgenic pig expressing *INS-SNAP* and demonstrate its suitability for *in vivo* labeling. We provide a strategy to overcome two major limitations of β-cell research: the translational gap between rodents and humans and *ex vivo* experiments for insulin turnover. We generated a transgenic pig as a model system since it closes this translational gap. Pigs are akin to humans in the anatomy of their gastrointestinal tract and the morphology and function of the pancreas (Wolf et al., 2014). Furthermore, it has been demonstrated that the structure and composition of pig pancreatic islets is much closer to human than to rodent islets (Hoang et al., 2014). In addition, molecular and developmental signature of islets and β-cells of pigs are more similar to that of humans, whereas more differences exist between that of rodents versus humans (Kim et al., 2020). This makes the pig a compelling model organism not only for the study of metabolism but also for islet and β-cell function. The age of insulin SGs has in the past been associated with the likelihood of exocytosis and recently a connection of the dysregulation of SG age with metabolic stress has been made (Yau et al., 2020). We chose the SNAP-tag fused with insulin in our pig model since it allows for conditional and flexible labeling of age-defined insulin SG pools with fluorophores of different colors over longer time spans compared to fluorescent timer proteins (Duncan et al., 2003). We used this technology successfully in isolated mouse islets to map age-distinct SG pools (Müller et al., 2017b). However, this knock-in mouse showed signs of impaired glucose tolerance (Ivanova et al., 2013), a problem that did not occur in the transgenic *SOFIA* pig. The major part of our knowledge on insulin SG trafficking is derived from *ex vivo* experiments using isolated islets or *in vitro* work with insulin-producing cell lines. Proteins tagged with self-labeling enzymes have so far been successfully used for *in vivo* studies in the mouse brain and in small animals (Masch et al., 2018; Poc et al., 2020; Campos et al., 2011; Yang et al., 2015) where application of the substrates is achieved by injection in the tissue. Administration of a single-color SNAP- or Halo-tag substrate via tail-vein injection has resulted in specific labeling in mice (Ivanova et al., 2013; Chen et al., 2021). This approach is less invasive and overcomes the substantial hurdles and potential severe complications, e.g. pancreatitis, associated with injections into a retroperitoneal and sensitive organ as the pancreas. Nevertheless, the high specificity of the SNAP substrates to their tag and the covalent binding upon contact allowed for specific labeling of insulin SGs even in adult pigs. We could demonstrate that a dual-color sequential labeling with only a relatively short time interval between the labeling steps results in partially distinct insulin SG pools. Future studies will aim to optimize substrate distribution to the β-cells as well as better control of the age-defined labeling. Ultimately, the *SOFIA* pig may be crossed to diabetic pig models (Renner et al., 2013, 2010) to gain insights into the changes in insulin SG turnover in T2D.

## ACKNOWLEDGEMENTS

The authors thank Christina Blechinger, Eva Jemiller, Anne Richter, Tatiana Schroeter (all LMU Munich, Germany) as well as Carla Münster, Daniela Friedland, Eyke Schöniger, Katharina Gan×, Carolin Wegbrod (all Paul Langerhans Institute Dresden, Germany), Cordula Andreé (Max Planck Institute for Cell Biology and Genetics, Dresden, Germany) and Bettina Mathes (Max Planck Institute for Medical Research, Heidelberg, Germany) for their excellent technical support. We thank Susanne Kretschmar (Center for Molecular and Cellular Bioengineering, Dresden, Germany) for electron microscopy processing. We are further grateful to Katja Pfriem for administrative assistance. This work was supported by the Light Microscopy Facility, a Core Facility of the CMCB Technology Platform at TU Dresden. This work was supported by the grant (82DZD00802) from the German Federal Ministry of Education and Research (BMBF) to the German Center for Diabetes Research (DZD) to E. Wolf and in part by the German Research Council (DFG; TRR127) to E. Wolf and E. Kemter. M. Solimena received funding from the German Center for Diabetes Research (DZD) by the German Ministry for Education and Research (BMBF), from the German Research Foundation (DFG) jointly with the Agence Nationale de la Recherche (grant SO 818/6-1), and from the Innovative Medicines Initiative 2 Joint Undertaking under grant agreements 115881 (RHAPSODY), which includes financial contributions from the European Union’s Framework Program Horizon 2020, EFPIA, and the Swiss State Secretariat for Education, Research and Innovation under contract 16.0097, as well as JDRF International and the Leona M. and Harry B. Helmsley Charitable Trust. A. Ivanova and A. Müller were the recipients of MeDDrive grants (60.372, 60.295, 60417) from the Carl Gustav Carus Faculty of Medicine at TU Dresden. M. Neukam was the recipient of a predoctoral fellowship from the Dresden International Graduate School for Biomedicine and Bioengineering (DIGS-BB) in the context of the Excellence Initiative of the German Research Foundation (DFG). T. Kurth and the EM facility of the Center for Molecular and Cellular Bioengineering are supported by the European Fund for Regional Development.

The authors declare no competing financial interests.

## Author contributions

Conceptualization: E. Kemter, A. Müller, A. Ivanova, M.Solimena, E. Wolf Investigation: E. Kemter, A. Müller, M. Neukam, A. Ivanova, N.Klymiuk, S. Renner, K. Yang, M. Kurome, V. Zakhartchenko, B. Kessler, K.P. Knoch, B. Ludwig, M. Bickle, H. Lickert, T. Kurth Resources: J. Broichhagen, K. Johnsson Formal analysis: E. Kemter, S. Renner, A. Müller, M. Neukam Writing-Original draft: E. Kemter, A. Müller, M. Solimena, E. Wolf Writing-Review and editing: E. Kemter, A. Müller, M. Neukam, J. Broichhagen, T. Kurth, M. Solimena, E. Wolf Visualization: A. Müller Supervision: E. Wolf, M. Solimena Funding acquisition: E. Kemter, A. Müller, M. Neukam, A. Ivanova, M. Solimena, E. Wolf

## Materials and Methods

### Generation of *SOFIA* pig

All animal procedures in this study were approved by the responsible animal welfare authority (Regierung von Oberbayern) and were performed according to the German Animal Welfare Act and Directive 2010/63/EU on the protection of animals used for scientific purposes. The SNAP-tag sequence (NEB) was cloned into a *β*-cell-specific expression vector with a porcine *INS* gene promoter, exon and intron sequence (Klymiuk et al., 2012) leading to a SNAP-tagged insulin protein product. The *INS-SNAP* vector was combined with a floxed neomycin resistance cassette. Porcine kidney cells were transfected with the linearized and excised expression vector, and pools of stable transfected male porcine kidney cell clones were generated. For Cre-mediated removal of floxed neo cassette, kidney cells were lipofected with a CAG-Cre expression vector directly before used for somatic cell nuclear transfer (SCNT) (Kurome et al., 2015). Cloned embryos were laparoscopically transferred to recipient gilts. Genotyping of offspring was performed by PCR using following primers: SNAP_1_for (5’-ACC AGA GCC ACT GAT GCA G -3’), SNAP_3_rev (5’-GGA GTG GCA CCT TCC AG -3’), *δ*neo_2_for (5’-CCT ACT TTC ACC AGC CTG AG -3’), *δ*neo_1_rev (5’-AGC TTG ATA TCG AAT TCC TGC AG -3’), neo_1_for (5’-ACA ACA GAC AAT CGG CTG CTC TG -3’) and neo_2_rev (5’-TGC TCT TCG TCC AGA TCA TCC TG -3’). Transgene integration patterns were analyzed by Southern blot analysis as described previously (Klose et al., 2005). Genomic DNA was extracted from skin using Wizard® Genomic DNA Purification Kit (Promega). 10μg DNA each were digested with *NdeI* or *Psh*AI, fractionated on 0.7% TBE agarose gel, and blotted under neutral conditions to Hybond™-XL nylon membrane (GE Healthcare). Probes were synthesized by PCR reaction from DNA of a transgenic animal using primers INS-for (5’-TCG TTA AGA CTC TAA TGA CCT C -3’) and INS-SNAP_5_rev (5’-ATC CCA GTT GCA GTA GTT CTC CAG C -3’) comprising the 3’-region of the porcine INS promoter and the 5’-region of the *INS-SNAP* sequence, microdialysed and ^32^P-labeled using Prime-a-Gene® Labeling System (Promega). Filters were prehybridized for 2 hr at 65°C in Rapid-hyb solution (GE Healthcare): Hybridization was performed overnight in the same buffer containing a ^32^P-labeled probe. After washing the membranes (once for 20 min at room temperature in 2x standard saline citrate containing 0.1% sodium dodecyl sulfate (SDS) and twice for 15 min at 65°C in 1x standard saline citrate containing 0.1% SDS), exposition of membranes were done in a Phosphor-Imager cassette overnight. Imaging plates were scanned with a Phosphor-Imager (Typhoon FLA9000; GE Healthcare).

### TLA sequencing

Commercially available targeted locus amplification (TLA) with subsequent next generation sequencing was performed by Cergentis, Utrecht, Netherlands.

### FACS sorting of *β*-cells and Western Blot

For FACS sorting of GFP-positive *β*-cells, the *SOFIA* pig was crossed with the *INS-eGFP* pig line expressing eGFP specifically in their *β*-cells (Kemter et al., 2017). Dual transgenic (*SOFIA/INS-eGFP*) and single *INS-eGFP* transgenic pigs were sacrificed at an age of 8-10 weeks, their pancreata were scissored, collagenase digested, sieved through 500 micron mesh and washed as described in (Kemter et al., 2017). Afterwards, single cell suspension by TrypLE™ Express enzyme digestion was prepared for FACS sorting as described elsewhere (Böttcher et al., 2021). Between 22,109 and 26,665 FACS-sorted *β*-cells were then lysed in Laemmli buffer and loaded on tricine gels for detection of insulin (Sigma, I2018), SNAP tag (NEB, P9310S) or GFP (MPI-CBG, Dresden, Germany, selfmade).

### Immunohistochemistry and confocal microscopy

Fixed pancreatic tissue was embedded in TissueTek, snapfrozen, and stored at −80°C for cryosectioning. Immunolabeling on cryosections was performed with anti-insulin (Sigma) and anti-SNAP (NEB) antibodies. Labeled cryo sections were imaged with a a Zeiss LSM 780 (DFG FZT 111) of the Light Microscopy Facility, a Core Facility of the CMCB Technology Platform at TU Dresden and a Nikon C2+ confocal microscope with 20x air and 40x and 63x oil immersion objectives.

### Transmission electron microscopy

Pancreatic tissue was fixed with 2.5% glutaraldehyde and 4% paraformaldehyde in 0.1 M Sörensen’s phosphate buffer (pH 7.4) at room temperature. After dehydration in a graded series of ethanol the specimens were embedded in epoxy resin as described before (Fava et al., 2012). Ultrathin sections were cut with a Leica EM UC6 ultramicrotome. After post-contrasting with uranyl-acetate and lead-citrate the sections were observed with an FEI Morgagni electron microscope running at 80 kV.

### Correlative light and electron microscopy (CLEM)

For CLEM pancreatic tissue was fixed with 2.5% glutaraldehyde and 4% paraformaldehyde in 0.1 M Sörensen’s phosphate buffer (pH 7.4). The specimens were processed for CLEM as described in (Völkner et al., 2021). In brief, they were embedded in 12% gelatine and immersed in 2.3 M sucrose at 4°C overnight. The samples were mounted on metal pins and immersed in liquid nitrogen. Ultrathin Tokuyasu sections were cut with a Leica EM UC6 ultramicrotome equipped with an FC6 cryo-unit. Sections were stained with the anti-SNAP antibody, followed by a AlexaFluor-conjugated secondary antibody. For immunogold-labelling protein-A-Gold 10 nm was applied followed by DAPI. Fluorescence imaging was done prior to electron microscopy with a Keyence Biozero 8000 fluorescence microscope as described in (Fabig et al., 2012).

### Intravenous glucose tolerance test (IVGTT)

An intravenous glucose tolerance test (IVGTT) was performed in 15-16-week-old *SOFIA* and *WT* control pigs. For stress-free, frequent blood sampling, central venous catheters (Careflow® 3 Fr, 200 mm, Merit Medical Systems) were surgically inserted into the external jugular vein via the vena auricularis under anesthesia as described previously (Renner et al., 2018). After an 18-hour fasting period, a bolus injection of concentrated 50% glucose solution (0.5 g per kg body weight) was administered through the central venous catheter. Blood samples were collected at the indicated time points (Fig. 3). Plasma glucose levels were determined using an AU480 autoanalyzer (Beckman-Coulter) and adapted reagent kits from Beckman-Coulter. Plasma insulin levels were determined by radioimmunoassay (Millipore). The net glucose elimination rate after glucose injection was calculated as the slope for the interval 1-30 minutes after glucose injection of the logarithmic transformation of the individual plasma glucose values. Insulin sensitivity indices were calculated according to Matsuda (ISI Matsuda) (Radikova, 2003). Acute insulin responses (AIR) were calculated as the difference of mean insulin levels at 1, 3, and 5 minutes following intravenous glucose load and basal insulin levels. Longitudinal data (glucose/insulin values during IVGTT) were statistically evaluated by analysis of variance (Linear Mixed Models; PROC MIXED; SAS 8.2), taking the fixed effects of Genotype (GT; transgenic, control), Time (relative to glucose application) and the interaction GT*Time as well as the random effect of the individual animal into account. AUC insulin/glucose was calculated using GraphPad Prism® software (version 5.02). AUCs and indices were tested for significance by Mann-Whitney-U-test using SPSS (version 21) software.

### *in vivo* labeling of Insulin-SNAP in pigs with SNAP substrates for imaging studies

BG-TMR and BG-SiR were synthesized as described previously (Keppler et al., 2003; Lukinavičius et al., 2013). Central venous catheters (Cavafix® Certo®, B. Braun) were surgically inserted into the external jugular vein via the *vena auricularis* at least one day before SNAP substrate or vehicle application. SNAP substrates were resuspended at a concentration of 1 μmol per 50 μl DMSO, and diluted directly prior to intravenous injection with a 50-fold volume of 0.9% NaCl. For single SNAP substrate *in vivo* labeling, 0.6 μmol BG-TMR per 100 kg BW or 1.8 - 2.0 μmol TMR-Star per 100 kg BW were intravenously injected in overnight fasted three *SOFIA* pigs and one *WT* pig. 5 hours after BG-TMR injection, autopsy took place for sampling of pancreas for histological analyses. For dual SNAP substrate *in vivo* labeling in two *SOFIA* pigs, 1.8 - 2.0 μmol per 100 kg BW BG-TMR as first SNAP substrate were intravenously injected at 5:15 pm during the last meal. The second SNAP substrate intravenous injection using BG-SiR at an amount of 1.8 - 1.9 μmol per 100 kg BW were done in overnight fasted animals at 8:15 am 15 hours after first SNAP substrate application. 2 hours after BG-SiR injection, autopsy took place. As a negative control to demonstrate specificity of SNAP substrate labeling of insulin-SNAP within insulin SGs in β-cells, one *SOFIA* pig received DMSO without substrate and one *WT* pig received BG-TMR substrate as described above.

## Supplementary Note 1: Supplementary Figures

**Supplementary Figure 1.**
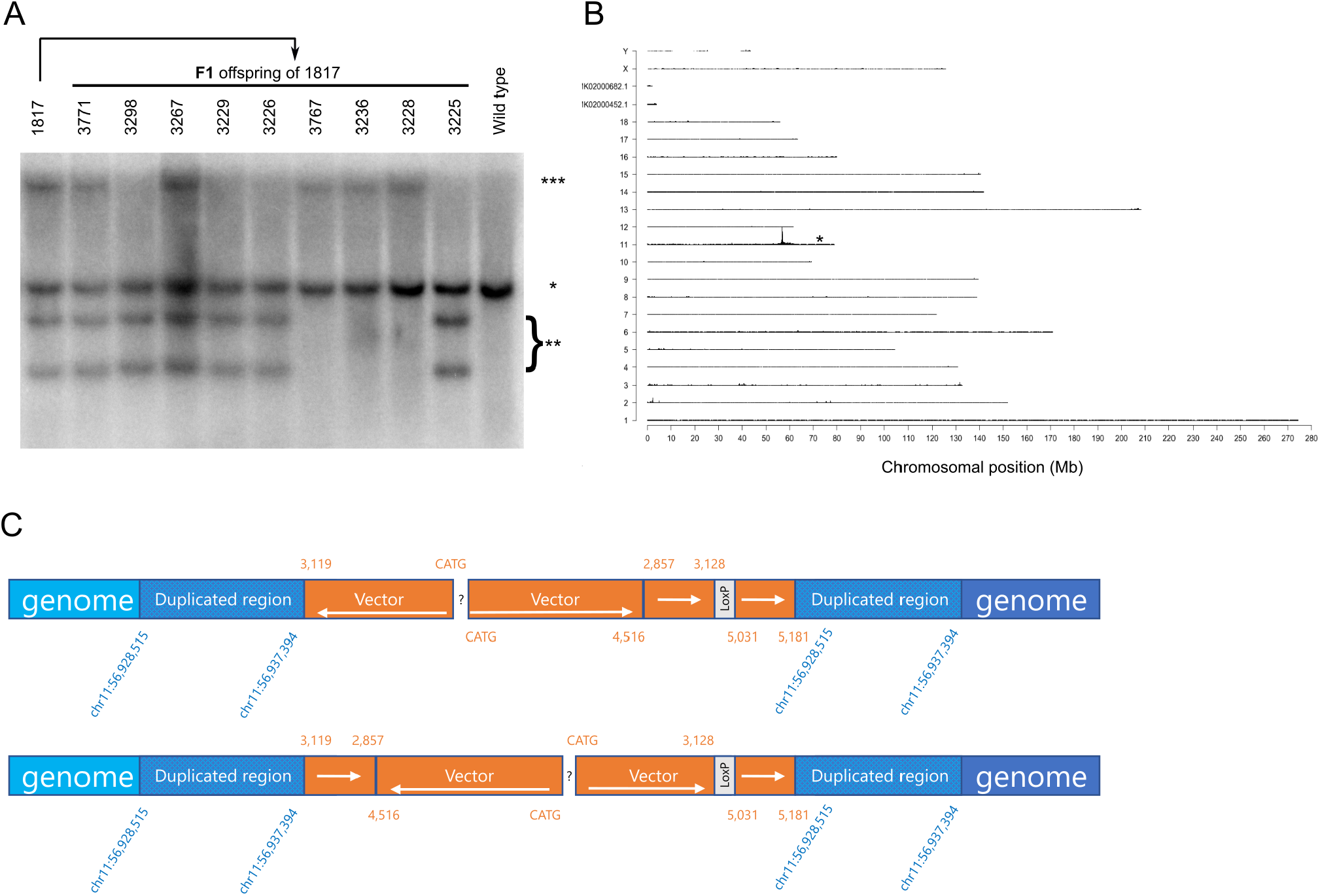
Southern blot and sequencing results of *SOFIA* pig offspring. (A) Southern blot analysis for validation of integration pattern of F1-offspring of founder 1817. DNA was digested with the ‘single cutter’ restriction endonuclease enzyme PshAI. *, the band representing the endogenous *INS* promoter; **, neo/δneo transgene integration site with two integrants where neo cassette was deleted only in one *INS-SNAP* integrant; ***, δneo transgene integrant. F1-offspring 3298, 3229, 3226 and 3225 possessed only neo/δneo transgene integration site, animals 3767, 3236 and 3228 had only δneo transgene integration site, whereas 3771 and 3267 harboured both transgene integration sites. (B) TLA sequencing of an animal from the *INS-SNAP* neo/δneo transgene integrant line revealed a single genomic transgene integration site in chromosome 11. (C) TLA sequencing derived interpretations of construct integration patterns into the identified single genomic transgene integration site.

**Supplementary Figure 2.**
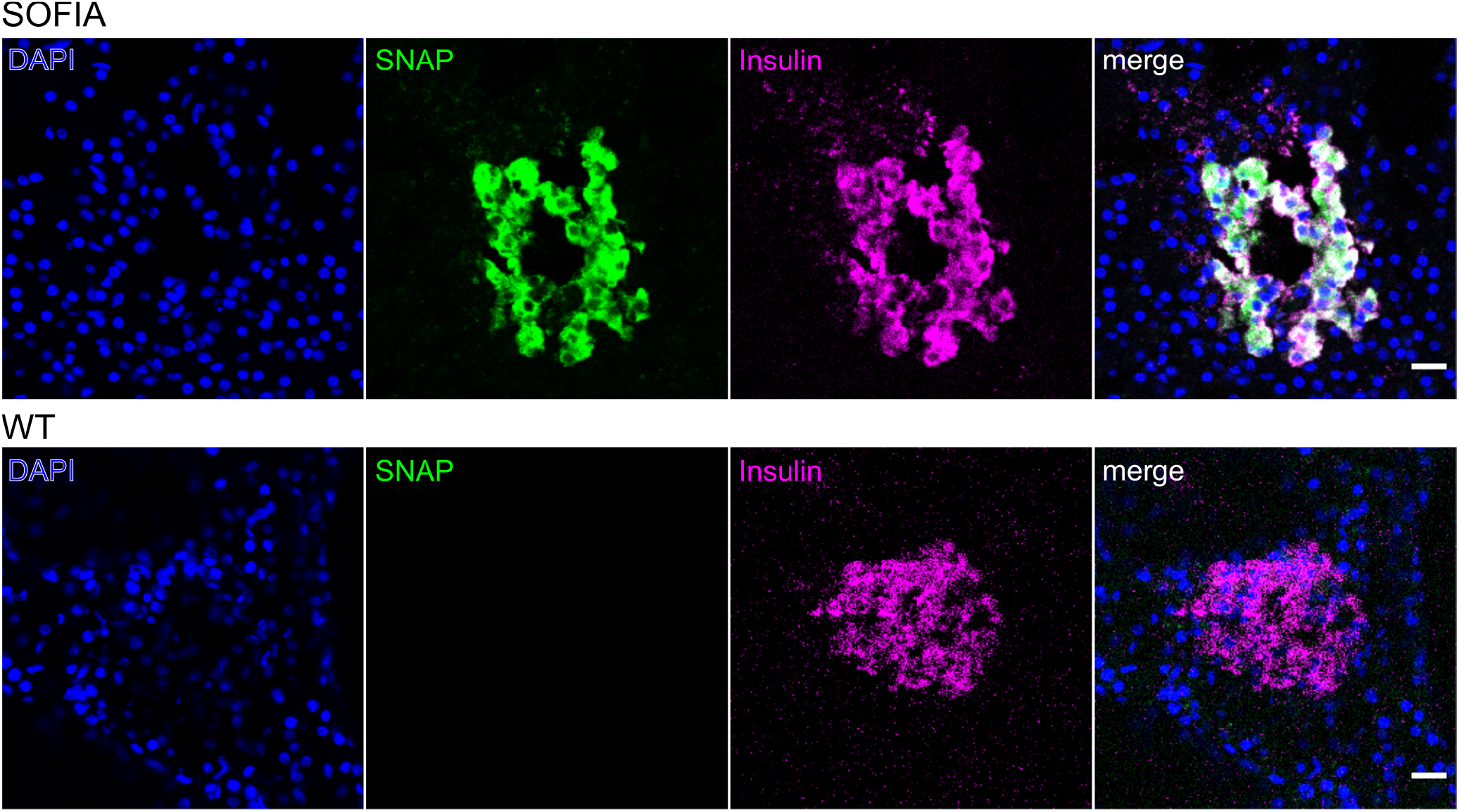
Immunolabeling against SNAP and insulin in *SOFIA* and *WT* pancreas. Scalebar: 20 μm

**Supplementary Figure 3.**
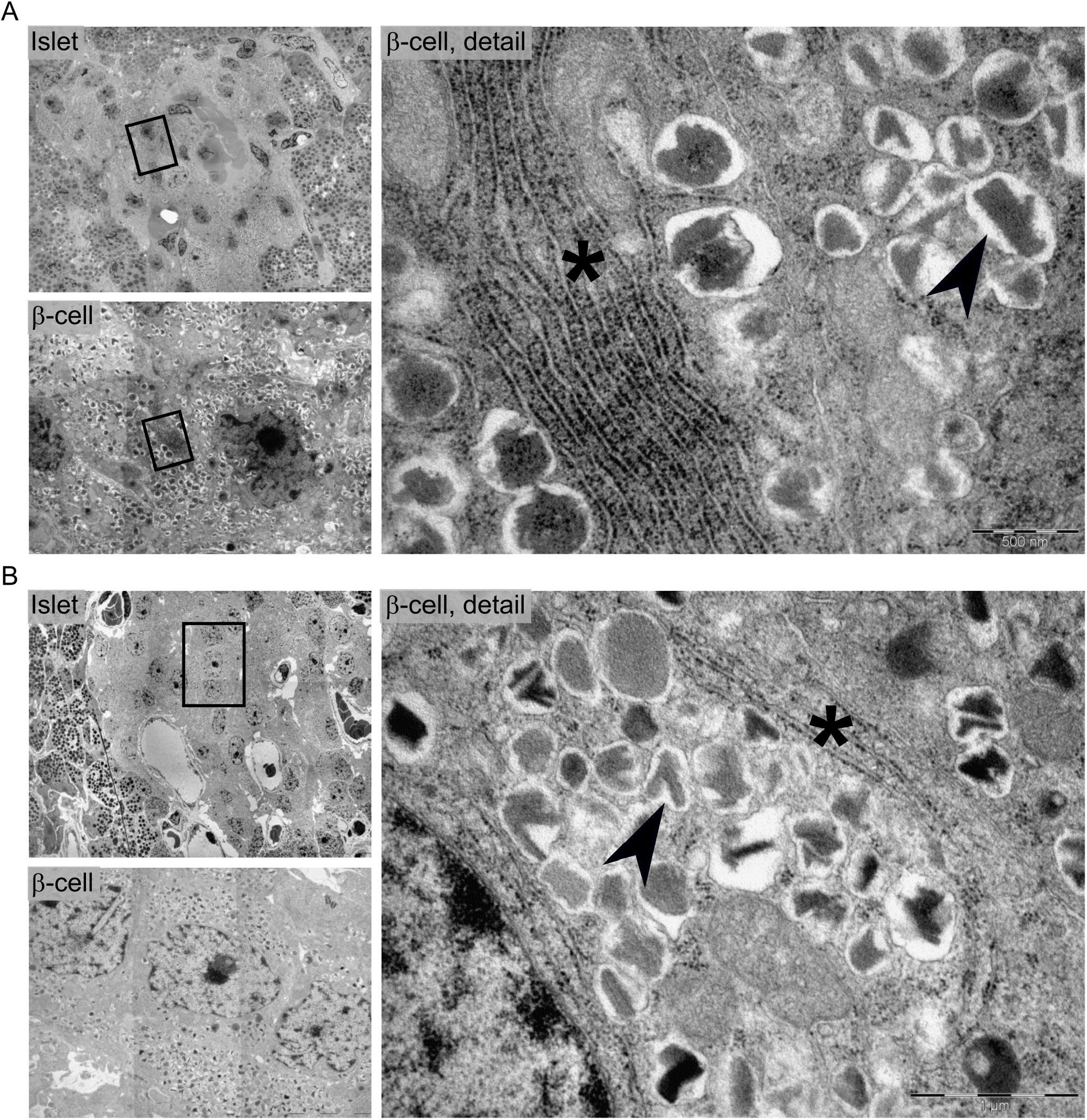
Comparison of *SOFIA* β-cell ultrastructure with that of pigs not expressing insulin-SNAP. A: Founder 1817 - *SOFIA* pig pancreas with SNAP B: Founder 1814 - pancreas of transgenic pig with neo cassette without SNAP expression

**Supplementary Figure 4.**
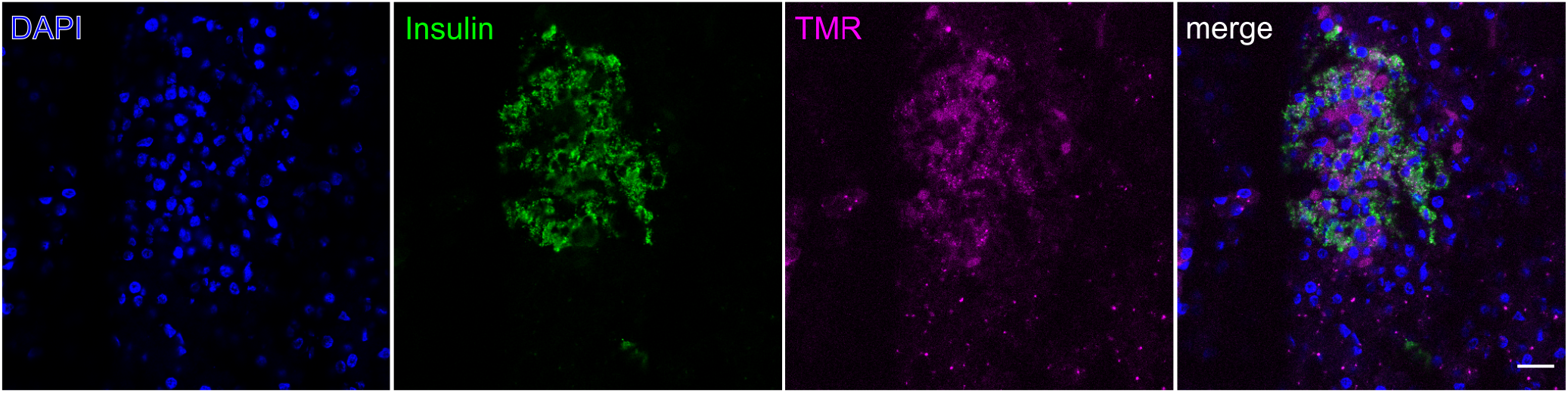
Immunolabeling against insulin on cryosections of *SOFIA* pig pancreas labeled with TMR. Scale bar: 20 μm

## References

S. Barg, L. Eliasson, E. Renström, and P. Rorsman. A Subset of 50 Secretory Granules in Close Contact With l-Type Ca2+ Channels Accounts for First-Phase Insulin Secretion in Mouse β-Cells. Diabetes, 51(suppl 1):S74–S82, Feb. 2002. ISSN 0012-1797, 1939-327X. doi: 10.2337/diabetes.51.2007.S74. URL https://diabetes.diabetesjournals.org/content/51/suppl_1/S74. Publisher: American Diabetes Association Section: Section 2: Biphasic Insulin Release: Pools and Signal Modulation.

A. Böttcher, M. Büttner, S. Tritschler, M. Sterr, A. Aliluev, L. Oppenländer, I. Burtscher, S. Sass, M. Irmler, J. Beckers, C. Ziegenhain, W. Enard, A. C. Schamberger, F. M. Verhamme, O. Eickelberg, F. J. Theis, and H. Lickert. Non-canonical Wnt/PCP signalling regulates intestinal stem cell lineage priming towards enteroendocrine and Paneth cell fates. Nature Cell Biology, 23(1):23–31, Jan. 2021. ISSN 1476-4679. doi: 10.1038/s41556-020-00617-2. URL https://www.nature.com/articles/s41556-020-00617-2. Number: 1 Publisher: Nature Publishing Group.

C. Campos, M. Kamiya, S. Banala, K. Johnsson, and M. González-Gaitán. Labelling cell structures and tracking cell lineage in zebrafish using SNAP-tag. Developmental Dynamics, 240(4):820–827, 2011. ISSN 1097-0177. doi: https://doi.org/10.1002/dvdy.22574. URL https://anatomypubs.onlinelibrary.wiley.com/doi/abs/10.1002/dvdy.22574. _eprint: https://anatomypubs.onlinelibrary.wiley.com/doi/pdf/10.1002/dvdy.22574.

E. Cerasi and R. Luft. The plasma insulin response to glucose infusion in healthy subjects and in diabetes mellitus. Acta Endocrinologica, 55(2):278–304, June 1967. ISSN 0001-5598. doi: 10.1530/acta.0.0550278.

S. Chen, Z. Huang, H. Kidd, M. Kim, E. H. Suh, S. Xie, E. H. Ghazvini Zadeh, Y. Xu, A. D. Sherry, P. E. Scherer, and W.-h. Li. In Vivo ZIMIR Imaging of Mouse Pancreatic Islet Cells Shows Oscillatory Insulin Secretion. Frontiers in Endocrinology, 12, 2021. ISSN 1664-2392. doi:10.3389/fendo.2021.613964. URL https://www.frontiersin.org/articles/10.3389/fendo.2021.613964/full?utm_source=S-TWT&utm_medium=SNET&utm_campaign=ECO_FENDO_XXXXXXXX_auto-dlvrit. Publisher: Frontiers.

C. M. Cohrs, J. K. Panzer, D. M. Drotar, S. J. Enos, N. Kipke, C. Chen, R. Bozsak, E. Schöniger, F. Ehehalt, M. Distler, A. Brennand, S. R. Bornstein, J. Weitz, M. Solimena, and S. Speier. Dysfunction of Persisting β Cells Is a Key Feature of Early Type 2 Diabetes Pathogenesis. Cell Reports, 31(1):107469, Apr. 2020. ISSN 2211-1247. doi: 10.1016/j.celrep.2020.03.033. URL https://www.sciencedirect.com/science/article/pii/S2211124720303478.

D. L. Curry, L. L. Bennett, and G. M. Grodsky. Dynamics of insulin secretion by the perfused rat pancreas. Endocrinology, 83(3):572–584, Sept. 1968. ISSN 0013-7227. doi: 10.1210/endo-83-3-572.

R. R. Duncan, J. Greaves, U. K. Wiegand, I. Matskevich, G. Bodammer, D. K. Apps, M. J. Shipston, and R. H. Chow. Functional and spatial segregation of secretory vesicle pools according to vesicle age. Nature, 422(6928):176–180, Mar. 2003. ISSN 1476-4687. doi:10.1038/nature01389. URL https://www.nature.com/articles/nature01389. Number: 6928 Publisher: Nature Publishing Group.

G. Fabig, S. Kretschmar, S. Weiche, D. Eberle, M. Ader, and T. Kurth. Labeling of ultrathin resin sections for correlative light and electron microscopy. Methods in Cell Biology, 111:75–93, 2012. ISSN 0091-679X. doi: 10.1016/B978-0-12-416026-2.00005-4.

E. Fava, J. Dehghany, J. Ouwendijk, A. Müller, A. Niederlein, P. Verkade, M. Meyer-Hermann, and M. Solimena. Novel standards in the measurement of rat insulin granules combining electron microscopy, high-content image analysis and in silico modelling. Diabetologia, 55(4): 1013–1023, Apr. 2012. ISSN 1432-0428. doi: 10.1007/s00125-011-2438-4.

P. A. Halban. Differential rates of release of newly synthesized and of stored insulin from pancreatic islets. Endocrinology, 110(4):1183–1188, Apr. 1982. ISSN 0013-7227. doi: 10.1210/endo-110-4-1183.

D.-T. Hoang, H. Matsunari, M. Nagaya, H. Nagashima, J. M. Millis, P. Witkowski, V. Periwal, M. Hara, and J. Jo. A Conserved Rule for Pancreatic Islet Organization. PLOS ONE, 9(10): e110384, Oct. 2014. ISSN 1932-6203. doi: 10.1371/journal.pone.0110384. URL https://journals.plos.org/plosone/article?id=10.1371/journal.pone.0110384. Publisher: Public Library of Science.

P. Hoboth, A. Müller, A. Ivanova, H. Mziaut, J. Dehghany, A. Sönmez, M. Lachnit, M. Meyer-Hermann, Y. Kalaidzidis, and M. Solimena. Aged insulin granules display reduced microtubule-dependent mobility and are disposed within actin-positive multigranular bodies. Proceedings of the National Academy of Sciences, 112(7):E667–E676, Feb. 2015. ISSN 0027-8424, 1091-6490. doi: 10.1073/pnas.1409542112. URL https://www.pnas.org/content/112/7/E667. Publisher: National Academy of Sciences Section: PNAS Plus.

A. Ivanova, Y. Kalaidzidis, R. Dirkx, M. Sarov, M. Gerlach, B. Schroth-Diez, A. Müller, Y. Liu, C. Andree, B. Mulligan, C. Münster, T. Kurth, M. Bickle, S. Speier, K. Anastassiadis, and M. Solimena. Age-Dependent Labeling and Imaging of Insulin Secretory Granules. Diabetes, 62(11):3687–3696, Nov. 2013. ISSN 0012-1797, 1939-327X. doi: 10.2337/db12-1819. URL https://diabetes.diabetesjournals.org/content/62/11/3687. Publisher: American Diabetes Association Section: Original Research.

M. Jaffredo, E. Bertin, A. Pirog, E. Puginier, J. Gaitan, S. Oucherif, F. Lebreton, D. Bosco, B. Catargi, D. Cattaert, S. Renaud, J. Lang, and M. Raoux. Dynamic Uni- and Multicellular Patterns Encode Biphasic Activity in Pancreatic Islets. Diabetes, 70(4):878–888, Apr. 2021. ISSN 0012-1797, 1939-327X. doi: 10.2337/db20-0214. URL https://diabetes.diabetesjournals.org/content/70/4/878. Publisher: American Diabetes Association Section: Islet Studies.

E. Kemter, C. M. Cohrs, M. Schäfer, M. Schuster, K. Steinmeyer, L. Wolf-van Buerck, A. Wolf, A. Wuensch, M. Kurome, B. Kessler, V. Zakhartchenko, M. Loehn, Y. Ivashchenko, J. Seissler, A. M. Schulte, S. Speier, and E. Wolf. INS-eGFP transgenic pigs: a novel reporter system for studying maturation, growth and vascularisation of neonatal islet-like cell clusters. Diabetologia, 60(6):1152–1156, June 2017. ISSN 1432-0428. doi: 10.1007/s00125-017-4250-2.

A. Keppler, S. Gendreizig, T. Gronemeyer, H. Pick, H. Vogel, and K. Johnsson. A general method for the covalent labeling of fusion proteins with small molecules in vivo. Nature Biotechnology, 21(1):86–89, Jan. 2003. ISSN 1087-0156. doi: 10.1038/nbt765.

S. Kim, R. L. Whitener, H. Peiris, X. Gu, C. A. Chang, J. Y. Lam, J. Camunas-Soler, I. Park, R. J. Bevacqua, K. Tellez, S. R. Quake, J. R. T. Lakey, R. Bottino, P. J. Ross, and S. K. Kim. Molecular and genetic regulation of pig pancreatic islet cell development. Development, 147(6), Mar. 2020. ISSN 0950-1991, 1477–9129. doi: 10.1242/dev.186213. URL https://dev.biologists.org/content/147/6/dev186213. Publisher: Oxford University Press for The Company of Biologists Limited Section: TECHNIQUES AND RESOURCES.

R. Klose, E. Kemter, T. Bedke, I. Bittmann, B. Kelsser, R. Endres, K. Pfeffer, R. Schwinzer, and E. Wolf. Expression of biologically active human TRAIL in transgenic pigs. Transplantation, 80(2):222–230, July 2005. ISSN 0041-1337. doi: 10.1097/01.tp.0000164817.59006.c2.

N. Klymiuk, L. v. Buerck, A. Bähr, M. Offers, B. Kessler, A. Wuensch, M. Kurome, M. Thormann, K. Lochner, H. Nagashima, N. Herbach, R. Wanke, J. Seissler, and E. Wolf. Xenografted Islet Cell Clusters From INSLEA29Y Transgenic Pigs Rescue Diabetes and Prevent Immune Rejection in Humanized Mice. Diabetes, 61(6):1527–1532, June 2012. ISSN 0012-1797, 1939-327X. doi: 10.2337/db11-1325. URL https://diabetes.diabetesjournals.org/content/61/6/1527. Publisher: American Diabetes Association Section: Immunology and Transplantation.

M. Kurome, B. Kessler, A. Wuensch, H. Nagashima, and E. Wolf. Nuclear Transfer and Transgenesis in the Pig. In N. Beaujean, H. Jammes, and A. Jouneau, editors, Nuclear Reprogramming: Methods and Protocols, Methods in Molecular Biology, pages 37–59. Springer, New York, NY, 2015. ISBN 978-1-4939-1594-1. doi: 10.1007/978-1-4939-1594-1_4. URL https://doi.org/10.1007/978-1-4939-1594-1_4.

G. Lukinavičius, K. Umezawa, N. Olivier, A. Honigmann, G. Yang, T. Plass, V. Mueller, L. Reymond, I. R. Corrêa Jr, Z.-G. Luo, C. Schultz, E. A. Lemke, P. Heppenstall, C. Eggeling, S. Manley, and K. Johnsson. A near-infrared fluorophore for live-cell super-resolution microscopy of cellular proteins. Nature Chemistry, 5(2):132–139, Feb. 2013. ISSN 1755-4349. doi: 10.1038/nchem.1546. URL https://www.nature.com/articles/nchem.1546. Number: 2 Publisher: Nature Publishing Group.

A. Lukinius and O. Korsgren. The Transplanted Fetal Endocrine Pancreas Undergoes an Inherent Sequential Differentiation Similar to That in the Native Pancreas: An Ultrastructural Study in the Pig-to-Mouse Model. Diabetes, 50(5):962–971, May 2001. ISSN 0012-1797,1939-327X. doi: 10.2337/diabetes.50.5.962. URL https://diabetes.diabetesjournals.org/content/50/5/962. Publisher: American Diabetes Association Section: Immunology and Transplantation.

J.-M. Masch, H. Steffens, J. Fischer, J. Engelhardt, J. Hubrich, J. Keller-Findeisen, E. D’Este, N. T. Urban, S. G. N. Grant, S. J. Sahl, D. Kamin, and S. W. Hell. Robust nanoscopy of a synaptic protein in living mice by organic-fluorophore labeling. Proceedings of the National Academy of Sciences, 115(34):E8047–E8056, Aug. 2018. ISSN 0027-8424, 1091-6490. doi: 10.1073/pnas.1807104115. URL https://www.pnas.org/content/115/34/E8047. Publisher: National Academy of Sciences Section: PNAS Plus.

A. Müller, H. Mziaut, M. Neukam, K.-P. Knoch, and M. Solimena. A 4D view on insulin secretory granule turnover in the β-cell. Diabetes, Obesity and Metabolism, 19(S1):107–114, 2017a. ISSN 1463-1326. doi: https://doi.org/10.1111/dom.13015. URL https://dom-pubs.onlinelibrary.wiley.com/doi/abs/10.1111/dom.13015. _eprint: https://dom-pubs.onlinelibrary.wiley.com/doi/pdf/10.1111/dom.13015.

A. Müller, M. Neukam, A. Ivanova, A. Sönmez, C. Münster, S. Kretschmar, Y. Kalaidzidis, T. Kurth, J.-M. Verbavatz, and M. Solimena. A Global Approach for Quantitative Super Resolution and Electron Microscopy on Cryo and Epoxy Sections Using Self-labeling Protein Tags. Scientific Reports, 7(1):23, Feb. 2017b. ISSN 2045-2322. doi: 10.1038/s41598-017-00033-x. URL https://www.nature.com/articles/s41598-017-00033-x. Number: 1 Publisher: Nature Publishing Group.

A. Müller, D. Schmidt, C. S. Xu, S. Pang, J. V. D’Costa, S. Kretschmar, C. Münster, T. Kurth, F. Jug, M. Weigert, H. F. Hess, and M. Solimena. 3D FIB-SEM reconstruction of micro-tubule–organelle interaction in whole primary mouse β cells. Journal of Cell Biology, 220 (e202010039), Dec. 2020. ISSN 0021-9525. doi: 10.1083/jcb.202010039. URL https://doi.org/10.1083/jcb.202010039.

M. Neukam, A. Sönmez, and M. Solimena. FLIM-based pH measurements reveal incretin-induced rejuvenation of aged insulin secretory granules. bioRxiv, page 174391, Nov. 2017. doi: 10.1101/174391. URL https://www.biorxiv.org/content/10.1101/174391v2. Publisher: Cold Spring Harbor Laboratory Section: New Results.

M. Ohara-Imaizumi, T. Fujiwara, Y. Nakamichi, T. Okamura, Y. Akimoto, J. Kawai, S. Matsushima, H. Kawakami, T. Watanabe, K. Akagawa, and S. Nagamatsu. Imaging analysis reveals mechanistic differences between first- and second-phase insulin exocytosis. The Journal of Cell Biology, 177(4):695–705, May 2007. ISSN 0021-9525. doi: 10.1083/jcb.200608132. URL https://www.ncbi.nlm.nih.gov/pmc/articles/PMC2064214/.

P. Poc, V. A. Gutzeit, J. Ast, J. Lee, B. J. Jones, E. D’Este, B. Mathes, M. Lehmann, D. J. Hodson, J. Levitz, and J. Broichhagen. Interrogating surface versus intracellular transmembrane receptor populations using cell-impermeable SNAP-tag substrates. Chemical Science, 11 (30):7871–7883, Aug. 2020. ISSN 2041-6539. doi: 10.1039/D0SC02794D. URL https://pubs.rsc.org/en/content/articlelanding/2020/sc/d0sc02794d. Publisher: The Royal Society of Chemistry.

Z. Radikova. Assessment of insulin sensitivity/resistance in epidemiological studies. Endocrine Regulations, 37(3):189–194, Sept. 2003. ISSN 1210-0668.

S. Renner, C. Fehlings, N. Herbach, A. Hofmann, D. C. v. Waldthausen, B. Kessler, K. Ulrichs, I. Chodnevskaja, V. Moskalenko, W. Amselgruber, B. Göke, A. Pfeifer, R. Wanke, and E. Wolf. Glucose Intolerance and Reduced Proliferation of Pancreatic β-Cells in Transgenic Pigs With Impaired Glucose-Dependent Insulinotropic Polypeptide Function. Diabetes, 59 (5):1228–1238, May 2010. ISSN 0012-1797, 1939-327X. doi: 10.2337/db09-0519. URL https://diabetes.diabetesjournals.org/content/59/5/1228. Publisher: American Diabetes Association Section: Original Article.

S. Renner, C. Braun-Reichhart, A. Blutke, N. Herbach, D. Emrich, E. Streckel, A. Wünsch, B. Kessler, M. Kurome, A. Bähr, N. Klymiuk, S. Krebs, O. Puk, H. Nagashima, J. Graw, H. Blum, R. Wanke, and E. Wolf. Permanent Neonatal Diabetes in INSC94Y Transgenic Pigs. Diabetes, 62(5):1505–1511, May 2013. ISSN 0012-1797, 1939-327X. doi: 10.2337/db12-1065. URL https://diabetes.diabetesjournals.org/content/62/5/1505. Publisher: American Diabetes Association Section: Original Research.

S. Renner, A. Blutke, B. Dobenecker, G. Dhom, T. D. Müller, B. Finan, C. Clemmensen, M. Bernau, I. Novak, B. Rathkolb, S. Senf, S. Zöls, M. Roth, A. Götz, S. M. Hofmann, M. Hrabě de Angelis, R. Wanke, E. Kienzle, A. M. Scholz, R. DiMarchi, M. Ritzmann, M. H. Tschöp, and E. Wolf. Metabolic syndrome and extensive adipose tissue inflammation in morbidly obese Göttingen minipigs. Molecular Metabolism, 16:180–190, Oct. 2018. ISSN 2212-8778. doi: 10.1016/j.molmet.2018.06.015.

P. Rorsman and E. Renström. Insulin granule dynamics in pancreatic beta cells. Diabetologia, 46 (8):1029–1045, Aug. 2003. ISSN 0012-186X. doi: 10.1007/s00125-003-1153-1.

P. Saeedi, I. Petersohn, P. Salpea, B. Malanda, S. Karuranga, N. Unwin, S. Colagiuri, L. Guariguata, A. A. Motala, K. Ogurtsova, J. E. Shaw, D. Bright, and R. Williams. Global and regional diabetes prevalence estimates for 2019 and projections for 2030 and 2045: Results from the International Diabetes Federation Diabetes Atlas, 9th edition. Diabetes Research and Clinical Practice, 157, Nov. 2019. ISSN 0168-8227, 1872-8227. doi: 10.1016/j.diabres.2019.107843. URL https://www.diabetesresearchclinicalpractice.com/article/S0168-8227(19)31230-6/abstract. Publisher: Elsevier.

H. Schatz, C. Nierle, and E. F. Pfeiffer. (Pro-)Insulin Biosynthesis and Release of Newly Synthesized (Pro-)Insulin from Isolated Islets of Rat Pancreas in the Presence of Amino Acids and Sulphonylureas+,++. European Journal of Clinical Investigation, 5(6):477–485, 1975. ISSN 1365-2362. doi: 10.1111/j.1365-2362.1975.tb02312.x. URL https://onlinelibrary.wiley.com/doi/abs/10.1111/j.1365-2362.1975.tb02312.x. _eprint: https://onlinelibrary.wiley.com/doi/pdf/10.1111/j.1365-2362.1975.tb02312.x.

M. Solimena, A. M. Schulte, L. Marselli, F. Ehehalt, D. Richter, M. Kleeberg, H. Mziaut, K.-P. Knoch, J. Parnis, M. Bugliani, A. Siddiq, A. Jörns, F. Burdet, R. Liechti, M. Suleiman, D. Margerie, F. Syed, M. Distler, R. Grützmann, E. Petretto, A. Moreno-Moral, C. Wegbrod, A. Sönmez, K. Pfriem, A. Friedrich, J. Meinel, C. B. Wollheim, G. B. Baretton, R. Scharfmann, E. Nogoceke, E. Bonifacio, D. Sturm, B. Meyer-Puttlitz, U. Boggi, H.-D. Saeger, F. Filipponi, M. Lesche, P. Meda, A. Dahl, L. Wigger, I. Xenarios, M. Falchi, B. Thorens, J. Weitz, K. Bokvist, S. Lenzen, G. A. Rutter, P. Froguel, M. von Bülow, M. Ibberson, and P. Marchetti. Systems biology of the IMIDIA biobank from organ donors and pancreatectomised patients defines a novel transcriptomic signature of islets from individuals with type 2 diabetes. Diabetologia, 61(3):641–657, Mar. 2018. ISSN 1432-0428. doi: 10.1007/s00125-017-4500-3. URL https://doi.org/10.1007/s00125-017-4500-3.

M. C. Vantyghem, J. Kerr-Conte, F. Pattou, M. H. Gevaert, C. Hober, A. Defossez, M. Mazzuca, and J. C. Beauvillain. Immunohistochemical and ultrastructural study of adult porcine endocrine pancreas during the different steps of islet isolation. Histochemistry and Cell Biology, 106(5):511–519, Nov. 1996. ISSN 1432-119X. doi: 10.1007/BF02473314. URL https://doi.org/10.1007/BF02473314.

M. Völkner, T. Kurth, L. Ebner, L. Bardtke, and M. O. Karl. Mouse retinal organoid growth and maintenance in longer-term culture. Frontiers in Cell and Developmental Biology, 9, 2021. ISSN 2296-634X. doi: 10.3389/fcell.2021.645704. URL https://www.frontiersin.org/articles/10.3389/fcell.2021.645704/abstract. Publisher: Frontiers.

E. Wolf, C. Braun-Reichhart, E. Streckel, and S. Renner. Genetically engineered pig models for diabetes research. Transgenic Research, 23(1):27–38, Feb. 2014. ISSN 1573-9368. doi: 10.1007/s11248-013-9755-y.

G. Yang, F. de Castro Reis, M. Sundukova, S. Pimpinella, A. Asaro, L. Castaldi, L. Batti, D. Bilbao, L. Reymond, K. Johnsson, and P. A. Heppenstall. Genetic targeting of chemical indicators in vivo. Nature Methods, 12(2):137–139, Feb. 2015. ISSN 1548-7091, 1548-7105. doi: 10.1038/nmeth.3207. URL http://www.nature.com/articles/nmeth.3207.

B. Yau, L. Hays, C. Liang, D. R. Laybutt, H. E. Thomas, J. E. Gunton, L. Williams, W. J. Hawthorne, P. Thorn, C. J. Rhodes, and M. A. Kebede. A fluorescent timer reporter enables sorting of insulin secretory granules by age. Journal of Biological Chemistry, 295(27):8901–8911, July 2020. ISSN 0021-9258, 1083-351X. doi: 10.1074/jbc.RA120.012432. URL https://www.jbc.org/article/S0021-9258(17)50315-4/abstract. Publisher: Elsevier.

